# Current population structure and pathogenicity patterns of *Ascochyta rabiei* in Australia

**DOI:** 10.1101/2020.12.21.423875

**Authors:** Ido Bar, Prabhakaran Thanjavur Sambasivam, Jenny Davidson, Lina M Farfan-Caceres, Robert C Lee, Kristy Hobson, Kevin Moore, Rebecca Ford

## Abstract

Ascochyta blight disease, caused by the necrotrophic fungus *Ascochyta rabiei*, is a major biotic constraint to chickpea production in Australia and worldwide. Detailed knowledge of the structure of the pathogen population and its potential to adapt to our farming practices is key to informing optimal management of the disease. This includes understanding the molecular diversity among isolates and the frequency and distribution of the isolates that have adapted to overcome host resistance across agro-geographically distinct regions.

Thanks to continuous monitoring efforts over the past six years, a comprehensive collection of *A. rabiei* isolates was collated from the major Australian production regions. To determine the molecular structure of the entire population, representative isolates from each collection year and growing region have been genetically characterised using a DArTseq™ genotyping-by-sequencing approach. The genotyped isolates were further phenotyped to determine their pathogenicity levels against a differential set of chickpea cultivars and genotype-phenotype associations were inferred.

Overall, the Australian *A. rabiei* population displayed a far lower genetic diversity (average Nei’s gene diversity of 0.047) than detected in other populations worldwide. This may be explained by the presence of a single mating-type in Australia, MAT1-2, limiting its reproduction to a clonal mode. Despite the low detected molecular diversity, clonal selection appears to have given rise to a subset of adapted isolates that are highly pathogenic on commonly employed resistance sources, and that are occurring at an increasing frequency.

To better understand the mechanisms and patterns of the pathogen adaptation, multi-locus genotype analysis was performed and two hypotheses were proposed on how new genotypes emerge. These were: 1) In a local, within-region evolutionary pathway; or 2) Through inter-region dispersal, most likely due to human activities. Furthermore, a cluster of genetically similar isolates was identified, with a higher proportion of highly aggressive isolates than in the general population, indicating the adaptive evolution of a sub-set of isolates that pose a greater risk to the chickpea industry.

The discovery of distinct genetic clusters associated with high and low isolate pathogenicity forms the foundation for the development of a molecular pathotyping tool for the Australian *A. rabiei* population. Application of such a tool, along with continuous monitoring of the genetic structure of the population will provide crucial information for the screening of breeding material and integrated disease management packages.

**Data Summary:** An online dataset containing all supporting genotyping and phenotyping data and the code required to reproduce the results, summary tables and plots found in this publication, is publicly available at Zenodo via the following links: https://zenodo.org/record/4311477; DOI: <u>10.5281/zenodo.4311477</u> (1).

## Introduction

With the ongoing growth of the human population, agricultural production systems are facing the increasing challenge of providing sustainable food sources to ensure food security for all (2). Particular emphasis is given to increasing the production of high nutrition plant-based protein sources, such as legumes, which in addition can fix nitrogen in the soil, making them ideal for crop rotation practices (3). Chickpea is one of the main crops of the legume family, known for high fibre, protein and vitamin content (4).

Australia is a major chickpea producer, with the bulk of the Australian crop primarily grown for export to the Indian sub-continent. With peak production exceeding 2 million tonnes in 2016 that was valued at over AU$2 billion, Australia is one of the world’s largest chickpea “cash crop” exporters (5). Chickpea is grown in Australia in several unique agro-geographical regions, mainly in the east of the continent, from central Queensland (QLD) through New South Wales (NSW), Victoria and South Australia (SA), with some minor production in western and northern Western Australia (WA) (Figure 1, reproduced from (6)). Each agro-geographical growing region is characterised by its distinct climate conditions, mainly temperature and rainfall, but also agronomic factors such as soil type as well as prevalence and risk of pests and pathogens, dictating which chickpea variety and farming practices are best suited for optimal production and yield (6–8).

**Figure 1.**
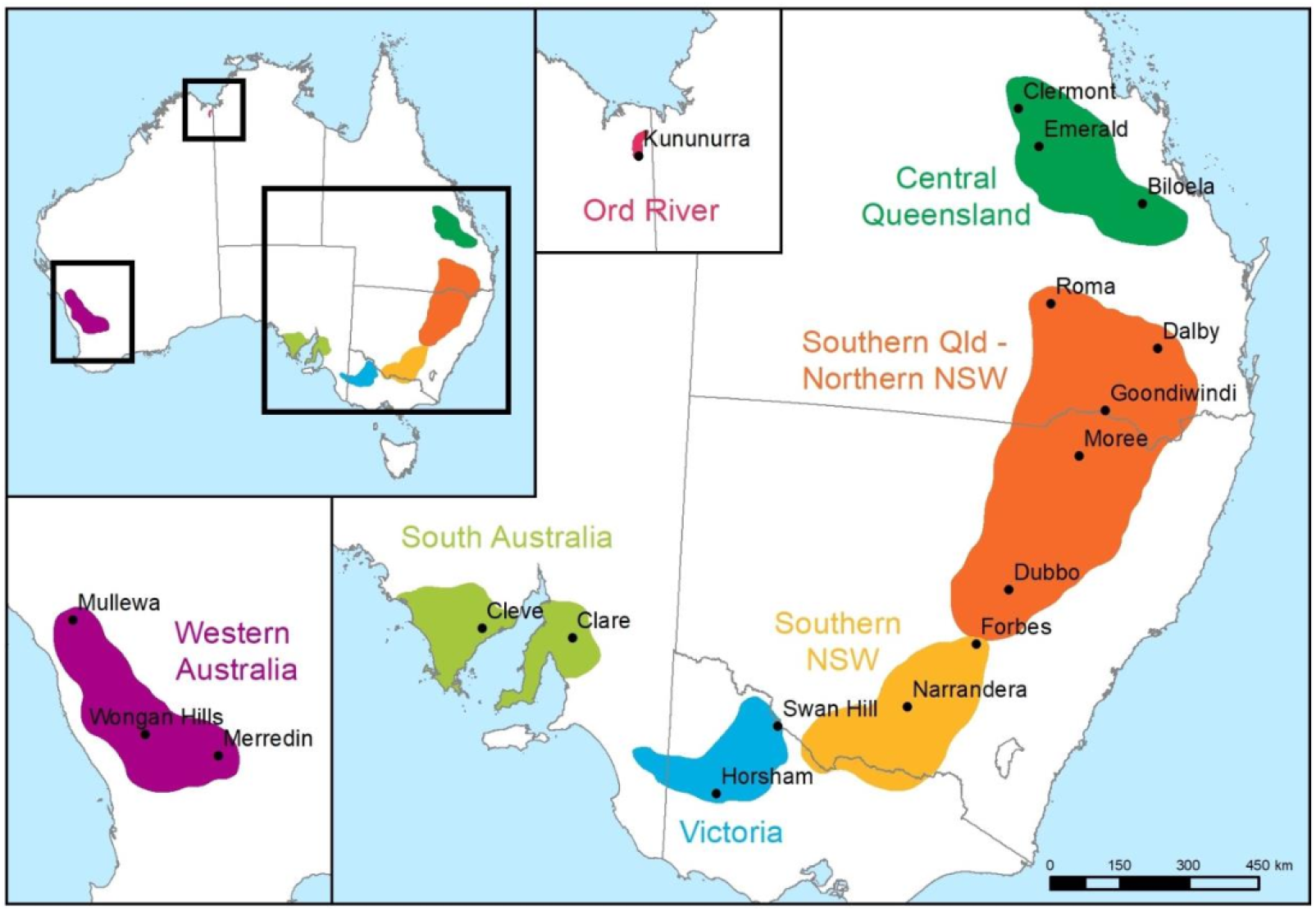
Chickpea growing regions in the states of Australia (reproduced with permission from (6))

Despite substantial expansion in Australian chickpea production over the past decade, from 230 kilotonnes (kt) in 2006-2007 to 2,000 kt in 2016-2017, yield remains threatened by a range of biotic (pests, weeds, pathogen-borne diseases) and abiotic (changes in climate conditions causing extreme temperatures, drought, frost or floods) factors.

One of the major biotic constraints to chickpea production is Ascochyta Blight, a disease caused by the necrotrophic fungal pathogen *Ascochyta rabiei*. Without proper informed disease management, which includes adoption of resistant cultivars, clean seed, pre-emptive fungicide spraying and rotation of crops, epidemics cause up to 100% yield loss (KM, personal communications). This translates to substantial economic losses to growers (9). The increased demand and record-high profits in Australia in 2016 led many farmers to overlook crop rotation and clean-seed practices, which might have contributed to a rise in the number of highly aggressive *A. rabiei* isolates, as was observed in the following years (10). These isolates are capable of causing severe disease symptoms, such as stem lesions and breakage on the most resistant chickpea cultivars (9,11). The mechanisms underlying this rapid change in pathogenicity are not yet understood.

To effectively manage the risk of Ascochyta blight outbreaks and to choose the most suitable cultivars for a growing region, as well as to advise on disease epidemiology, prevention and management strategies, it is essential to understand the pathogen population structure within each growing region across Australia. This will help to assess the adaptation potential of *A. rabiei* to overcome host resistance and chemical management strategies implemented by growers. Furthermore, the potential identification of fungal adaptation patterns unique to a growing region will assist in predicting future fungal evolution and epidemiology in response to particular farming practices and/or climatic factor trends.

The Australian *A. rabiei* population is thought to be reproducing largely clonally, with only a single mating-type, MAT1-2 detected in a population study held during 2009-2010 (12). This mode of reproduction explains the low genetic diversity observed in Australia relative to other populations worldwide, as was demonstrated by analysis of simple sequence repeat (SSR) markers (11–13). SSR markers are generally highly polymorphic, cost efficient to use and have been used extensively in population genetics studies (14). High mutation rates and assumed neutral evolution of SSR loci results in the accumulation of numerous population-specific alleles, revealing hidden population structures. However, the high allelic variability of SSR markers may also mask patterns of genomewide genetic diversity (15). Furthermore, in a population with low genetic diversity, such as the Australian *A. rabiei* population, a larger number of SSR markers is required to accurately distinguish between individuals and to assign haplotypes. The development, validation and application of a large number of informative SSR markers is labour intensive, time consuming and costly (16). Alternatively, a different approach for genotyping should be considered that is based on an alternative and more informative marker type.

Single nucleotide polymorphisms (SNPs) are genetic variants that occur every 100-300 bp on average across all regions of a genome, including within and in close proximity to gene coding regions. Moreover, heterozygosity estimates, such as Nei’s gene diversity (expected heterozygosity), can be more readily applied to bi-allelic SNPs than to multi-allelic SSRs to produce consistent estimates (15). Genotyping using high-throughput sequencing technologies, such as whole-genome sequencing (WGS) and genotyping-by-sequencing (GBS), allows an accurate identification of thousands of SNPs across entire or partial representations of genomes without prior knowledge of the genome (*de novo*). The relatively small size of the *A. rabiei* genome (~41 Mbp, <u>GCA 004011695.1</u> (17)), means that these methods are affordable to use with large sample sizes for population studies and even high-resolution genome-wide association studies (GWAS), to associate variants with traits of interest (18).

In this study, a GBS approach was employed to identify SNP markers among a representative collection of Australian *A. rabiei* isolates. The SNP data was then used to infer population structure within and among growing regions and to identify SNPs, and hence potential genes, associated with high levels of pathogenicity towards commonly grown chickpea cultivars.

## Methods

### *Ascochyta rabiei* isolates

The Australian *A. rabiei* isolates used in this study were collected between 2013 and 2018 inclusive, as part of a 6-year monitoring program funded by the Grain Research and Development Corporation (GRDC project ID UM00052; https://grdc.com.au/research/projects/project?id=2023). Another two isolates P2 and P4 (19), provided by Prof. Diego Rubiales (Institute for Sustainable Agriculture, Cordoba, Spain) were derived from the International Center for Agricultural Research in the Dry Areas (ICARDA) collection. These two isolates were confirmed as MAT 1-1 using the PCR assay developed by Barve et al. (20) and were used as outgroups. All of the isolates were confirmed as *A. rabiei* through disease symptomology and species-specific molecular testing (20). Passport data for the isolates is provided in Supplementary Table 1 including source (geographic and host cultivar) and date of collection. Disease scoring assays were performed in a controlled-environment growth room using a mini-dome assay to classify the isolates into pathogenicity groups, as described by Chen *et al*. (21) and refined by Mehmood *et al*. (11) and Sambasivam *et al*. (22). Specifically, isolates were assessed for their ability to cause disease symptoms on a differential set of chickpea cultivars, ranging from resistant to susceptible in their reaction to Ascochyta blight (ICC3996, Genesis090, PBA HatTrick and Kyabra, respectively) (22). The disease reaction on each host differential was ranked as low, moderate or high based on a 1-9 qualitative scale (21,23) and the overall pathogenicity group (PG) was assigned by the additive effect, with specific criteria as detailed in Table 1.

**Table 1.** An example of pathogenicity group assignment of *A. rabiei* isolates based on disease responses on the differential host set

| Isolate | Year | State | ICC3996 | Genesis090 | HatTrick | Kyabra | Patho. Group<sup>a</sup> |
| --- | --- | --- | --- | --- | --- | --- | --- |
| 17CUR005 | 2017 | VIC | High | High | High | High | 5 |
| FT13092-1 | 2013 | SA | Moderate | High | High | High | 4 |
| F15027 | 2015 | SA | Moderate | Moderate | High | High | 3 |
| 16CUR014 | 2016 | VIC | Low | Moderate | High | High | 2 |
| AS18084 | 2018 | VIC | Moderate | Low | High | High | 2 |
| TR6400 | 2014 | NSW | Low | Low | High | High | 1 |
| 13RUP002 | 2013 | VIC | Low | Low | Low | High | 0 |

<sup>a</sup>Classification into Pathogenicity Groups (PG) was determined by the following criteria: Low disease response on all hosts – PG0; High on PBA HatTrick and low on ICC3996 and Genesis090 – PG1; High on PBA HatTrick and a combination of Low and Moderate on ICC3996 and Genesis090 – PG2; High on PBA HatTrick and Moderate on both ICC3996 and Genesis090 – PG3; High on PBA HatTrick and Genesis090 and Moderate on ICC3996 – PG4; and High on ICC3996, Genesis090 and PBA HatTrick – PG5.

### Genotyping-by-Sequencing

GBS genotyping was performed on the two outgroup isolates sourced from ICARDA and a subset of 279 isolates that was selected from the Australian *A. rabiei* collection described above, representing the range of collection years, geographical regions, host cultivars and pathogenicity groups (see details in Supplementary Table S1). Numbers of isolates assessed from each of the classification criteria varied and were dependent on the occurrence of an epidemic within a region and within a year (Table 2). DNA was extracted from the selected 279 isolates using a modified CTAB extraction method (24). The DNA samples were genotyped using DArTseq™ (Diversity Array Technologies P/L, Canberra, Australia) (25,26).

**Table 2.** Number of Australian *A. rabiei* isolates used for DArTseq™ genotyping by Year and State^a^

| Year | NSW | QLD | SA | VIC | WA | Total |
| --- | --- | --- | --- | --- | --- | --- |
| 2013 | 0 | 0 | 23 | 3 | 0 | 26 |
| 2014 | 24 | 0 | 4 | 8 | 0 | 36 |
| 2015 | 0 | 0 | 9 | 7 | 0 | 16 |
| 2016 | 8 | 5 | 12 | 5 | 13 | 43 |
| 2017 | 17 | 16 | 14 | 15 | 11 | 73 |
| 2018 | 16 | 1 | 24 | 20 | 24 | 85 |
| Total | 65 | 22 | 86 | 58 | 48 | **279** |

<sup>a</sup> Australian states are abbreviated in the table as follows: NSW, New South Wales; QLD, Queensland; SA, South Australia; VIC, Victoria; WA, Western Australia, as in Figure 1.

### Population genetics analysis

Variant tables produced by DArTseq™ were imported into the R language and environment for statistical computing as *adegenet* (v2.1.1) genlight objects (27) using the *dartR* (v1.1.11) package (27–29). The SNP dataset was filtered to remove loci that did not show any variability (monomorphic) or did not pass the filtering criteria of reproducibility (DArTseq™ RepAvg score >0.95) or coverage (read depth >5). Also, loci and individuals with low genotype call rates (>20% missing data) were removed. When multiple SNPs were identified on a single tag (locus), a single representative SNP was selected from each tag using the “best” approach implemented in *dartR*. The locus was removed if it contained five or more SNPs.

The filtered SNP set was used to calculate a matrix of pairwise genetic distances between each pair of isolates, which was used for principal components (PC) analysis with the *ade4* (v1.7.13) and *adegenet* R packages (29–32). This was achieved by retaining principal components that explained at least 60% of the cumulative variance. Summarised genetic similarity between populations were assessed from the SNP set by calculating the Euclidean distances and presented as a neighbour-joining phylogenetic tree, as implemented by the gl.tree.nj() function in *dartR* (28).

#### Clonality and population analysis

To assess the level of clonality in the population, the retained SNP markers were used to define multilocus genotypes (MLGs), representing a unique combination of alleles in each isolate, while allowing a minimal degree of freedom using a distance threshold ~1, to allow for genotyping errors and missing values. Isolates that shared the same MLG were considered to be clones of the same lineage and were collapsed together for downstream analysis of the *A. rabiei* population structure. To visualise the genetic relationships between MLGs, a minimum spanning network (MSN) was constructed, based on a Euclidean distance dissimilarity matrix between each pair of MLGs. Population diversity metrics, including Shannon-Weiner Diversity index; Stoddard and Taylor’s Index; Simpson’s index; Corrected Simpson’s index (Λ · *N*/(*N* – 1)); Genotypic Evenness index; Nei’s gene diversity (expected heterozygosity); and Clonal Fraction (1 – MLG/JV), were calculated using *poppr* (v2.8.3) based on the MLG analysis (33,34). The analyses were performed following the best practices for population genetic analysis for clonal fungal pathogens, as specified by Grünwald *et al*. (35).

#### Isolate relatedness

To determine the pairwise genetic distances between isolates, the Euclidean distance between each pair of isolates was calculated based on the allele frequencies of the filtered SNP set (36). The distance matrix was used by *pheatmap* (v1.0.12) (37) to cluster the isolates and plot a heatmap and a dendrogram to visually represent isolate relatedness and identify patterns associated with origin and pathogenicity. The proportion of MLGs comprising isolates from multiple clusters was calculated to assess the robustness of the MLG and cluster assignment.

Isolates were further classified into “low” and “high” pathogenicity if they were assigned to pathogenicity groups 0-2 or 3-5, respectively. This separated between isolates that were able to cause moderate and severe disease symptoms on the more resistant host cultivars Genesis 090 and ICC3996. This classification was converted into a binary format (0,1) and used as the ‘response’ in a generalized linear mixed-effect model (GLMM) to determine the effect of the assigned cluster, using a binomial model, as implemented by the *lme4* (v1.1.21) R package (38,39). The odds ratio was calculated to reflect the differences in proportions of “high” and “low” pathogenicity isolates between the clusters and the Tukey method was applied on the log of the odds ratio scale to determine statistical significance.

#### Variant association with pathogenicity

To identify putative allelic variants that contributed to pathogenicity, a genome-wide-association-study (GWAS) approach was employed using the filtered SNP data of the genotyped population. The analysis was performed with *SNPassoc* (v1.9.2) R package (40), assuming a “log-additive” model and a significance threshold that was adjusted to reduce the false discovery rate in the case of multiple testing (Q-value ≤ 0.05) (41).

Annotation of the genes at each significantly associated SNP locus was performed based on the reference genome and gene models of A. *rabiei* isolate ArME14 (17). Additional functional annotation of these loci was performed using homology searches using BLAST against the NCBI non-redundant protein database (nr) and by InterProScan against the InterPro conglomerate databases (42,43). Effector prediction of the variant-associated sequences was performed by EffectorP (v1.0/2.0) (44,45).

#### Supporting Data

An online dataset containing all supporting genotyping and phenotyping data and the code required to reproduce the results, summary tables and plots found in this publication, is publicly available at Zenodo via the following links: https://zenodo.org/record/4311477; DOI: <u>10.5281/zenodo.4311477</u> (1).

## Results

While the main focus of this study was the genetic diversity of the Australian *A. rabiei* population, we included two MAT 1-1 isolates from a Northern Hemisphere collection (ICARDA), distinct from the isolated population from Australia, in our genotyping analysis. A total of 1,474 single nucleotide polymorphisms (SNPs) were detected from all the samples and was used for PC analysis. The analysis revealed that most of the variance (approximately 82%) in the genetic distance between the isolates was derived from the two outgroup isolates sourced from ICARDA (Figure 2A). The Australian isolates showed very low genetic diversity and overlay each other in Figure 2A. The two outgroup isolates displayed a significantly greater genetic diversity compared to the Australian isolates and to one another. The genetic distance within the Australian isolates was visible only when focusing on the 3^rd^ and 4^th^ PC and showed a major cluster of very similar isolates from all regions, and three distinct clusters of isolates separated by their state of origin (Figure 2B). Nei’s gene diversity for the P2 and P4 isolates from ICARDA was 0.313, and that of the Australian and ICARDA isolates collectively as a single population was 0.102. Inclusion of P2 and P4 isolates from the ICARDA collection in a phylogenetic tree illustrated the potential extent of A. *rabiei* genetic diversity globally and emphasized the limited genetic diversity in the Australian population (Supplementary Figure S1).

**Figure 2.**
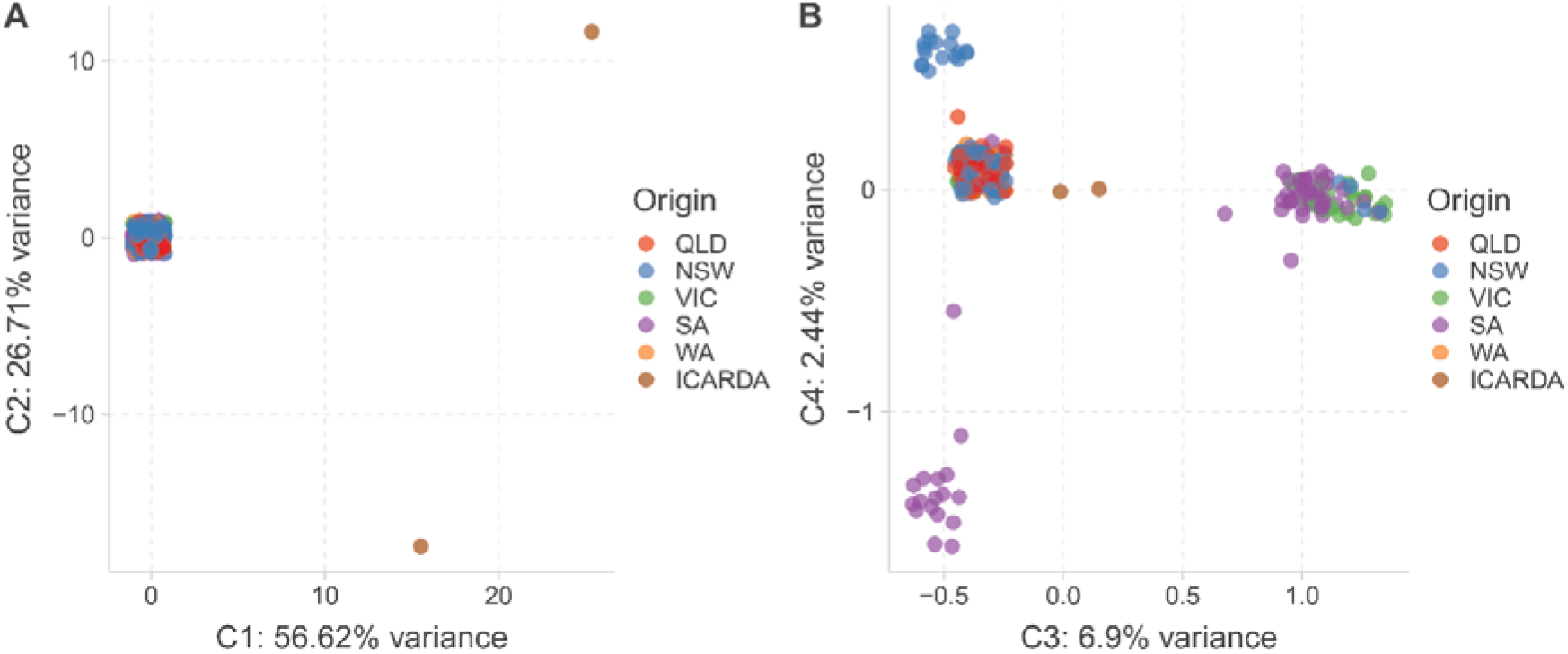
Principal component analysis representing the genetic distance between all isolates. The first two principal components are presented in **A** and components 3 and 4 are presented in **B**. Isolates are coloured by their origin: QLD - Queensland; NSW - New South Wales; VIC - Victoria; SA - South Australia; WA - Western Australia; and ICARDA – International Center for Agricultural Research in the Dry Areas (Middle East).

### The Australian *A. rabiei* population structure inferred from DArTseq^™^

After removal of the two outgroup isolates and focusing on the sampled population of Australian *A. rabiei* isolates (n=279), only 229 SNPs were polymorphic. Additional quality filtration removed one locus with a high rate of missing data (>20%), seven loci with low reproducibility scores (<0.95) and an additional nine secondary SNPs that shared the same sequence tags (loci) and were predicted to be linked to existing markers. Finally, 212 high-confidence polymorphic SNPs were retained for subsequent analyses of the Australian population.

The retained SNPs were used to define multilocus genotypes (MLGs), representing a unique combination of alleles for each isolate, in a similar way that the SSR haplotypes were previously reported and assessed for Australian *A. rabiei* (11). For this, 185 contracted MLGs were determined to represent the genotypic variability among the sampled 279 isolates, with 94 isolates considered as clones.

The Australian *A. rabiei* population contained very low genetic diversity across the years sampled with an average expected heterozygosity (Hexp) of 0.016 and minor seasonal fluctuations ranging from a low of Hexp = 0.012 in 2015 to Hexp = 0.021 in 2013. The observed higher diversity in 2013 was supported by a higher number of expected MLGs and low clonal fraction (Table 3). An examination of the population diversity within each state revealed an extremely low diversity among isolates originating from WA (Hexp= 0.006 and CF = 0.458), indicating that close to half of the 48 isolates (within our collection) were considered as genetic clones of existing MLGs. Higher diversity was found among isolates collected in other states, with similar Hexp of approximately 0.016±0.001, and particularly in QLD with a CF=0 (Table 3).

**Table 3.** *Ascochyta rabiei* population diversity metrics derived from DArTseq™ data within each collection year (2013-2018) and geographic region (state).

| Pop | N | MLG | eMLG | SE | H | G | Hexp | E.5 | Lambda | Lambda* | CF |
| --- | --- | --- | --- | --- | --- | --- | --- | --- | --- | --- | --- |
| 2013 | 26 | 25 | 15.631 | 0.483 | 3.205 | 24.143 | 0.021 | 0.979 | 0.959 | 0.997 | 0.038 |
| 2014 | 36 | 30 | 14.356 | 1.065 | 3.283 | 21.600 | 0.016 | 0.803 | 0.954 | 0.981 | 0.167 |
| 2015 | 16 | 12 | 12.000 | 0.000 | 2.361 | 9.143 | 0.012 | 0.848 | 0.891 | 0.950 | 0.250 |
| 2016 | 43 | 33 | 13.823 | 1.239 | 3.318 | 20.775 | 0.014 | 0.743 | 0.952 | 0.975 | 0.233 |
| 2017 | 73 | 54 | 14.136 | 1.257 | 3.756 | 29.442 | 0.015 | 0.681 | 0.966 | 0.979 | 0.260 |
| 2018 | 85 | 57 | 13.747 | 1.353 | 3.742 | 26.661 | 0.013 | 0.623 | 0.962 | 0.974 | 0.329 |
| QLD | 22 | 22 | 22.000 | 0.000 | 3.091 | 22.000 | 0.015 | 1.000 | 0.955 | 1.000 | 0.000 |
| NSW | 65 | 51 | 19.640 | 1.335 | 3.791 | 35.504 | 0.016 | 0.797 | 0.972 | 0.987 | 0.215 |
| VIC | 58 | 45 | 19.012 | 1.434 | 3.625 | 27.574 | 0.015 | 0.727 | 0.964 | 0.981 | 0.224 |
| SA | 86 | 64 | 18.824 | 1.571 | 3.883 | 28.667 | 0.018 | 0.582 | 0.965 | 0.976 | 0.256 |
| WA | 48 | 26 | 14.233 | 1.607 | 2.745 | 7.629 | 0.006 | 0.455 | 0.869 | 0.887 | 0.458 |
| **Total** | **279** | **185** | **19.191** | **1.633** | **4.738** | **42.236** | **0.016** | **0.364** | **0.976** | **0.980** | 0.337 |

To visualise the genetic relationships detected between MLGs, a minimum spanning network (MSN) was constructed, highlighting a number of isolates in each MLG and their state of origin (Figure 3).

**Figure 3.**
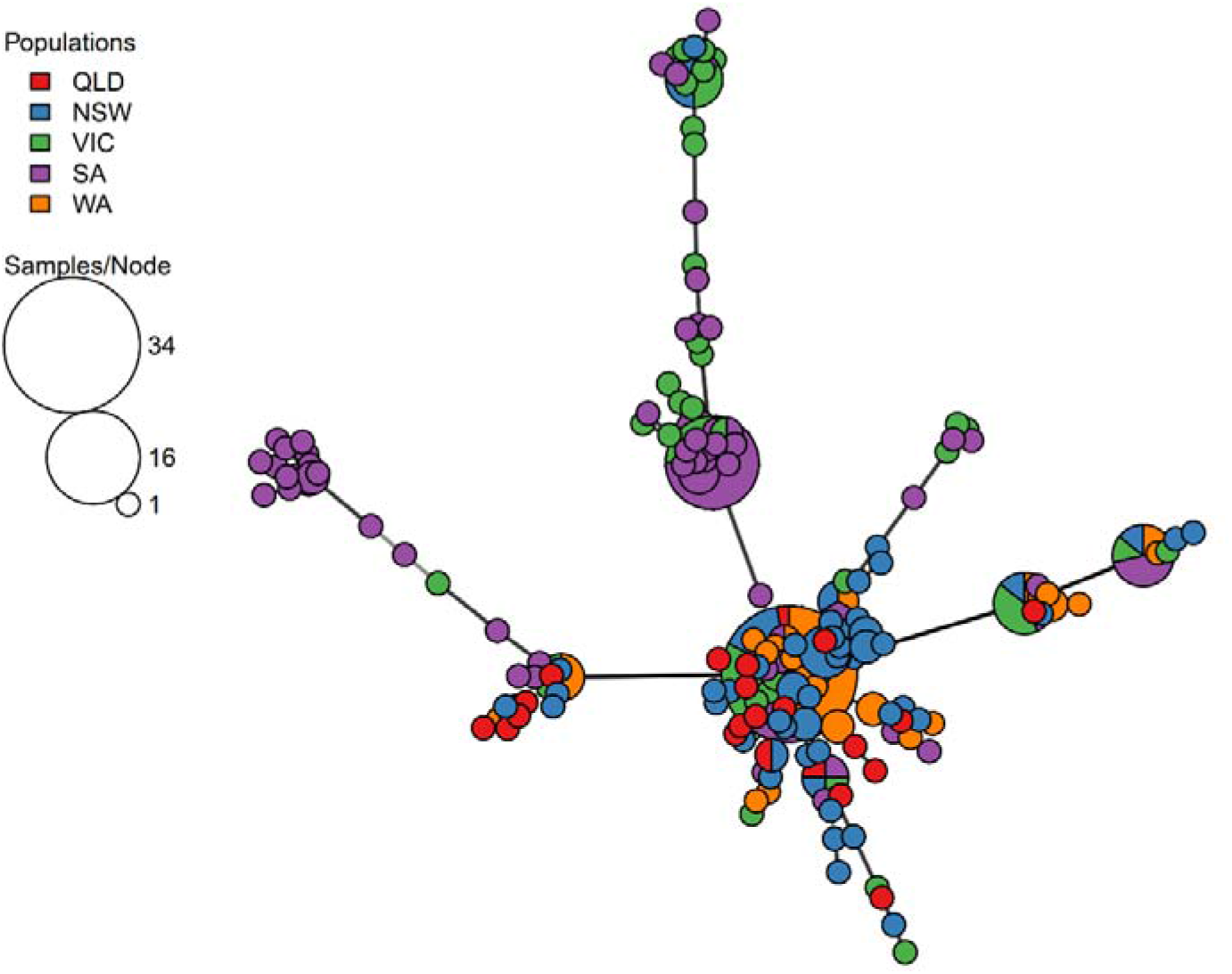
Minimum spanning network diagram of multilocus genotypes by state. Where node size represents the number of isolates identified in each MLG, and the length of the branches between the nodes represents the genetic distance. Node colours represent the origin of the isolates in the MLG.

A closer examination of several haplotypes highlighted differences in their occurrence, frequencies, and distributions among states. For example, isolates of MLG.264 were abundant in WA (11 isolates, comprising 39.3% of the isolates in the MLG) but these were also found in high frequencies in SA (8 isolates), and VIC (6 isolates), and to a lesser extent in NSW (2 isolates) and QLD (1 isolate) (Figure 4).

**Figure 4.**
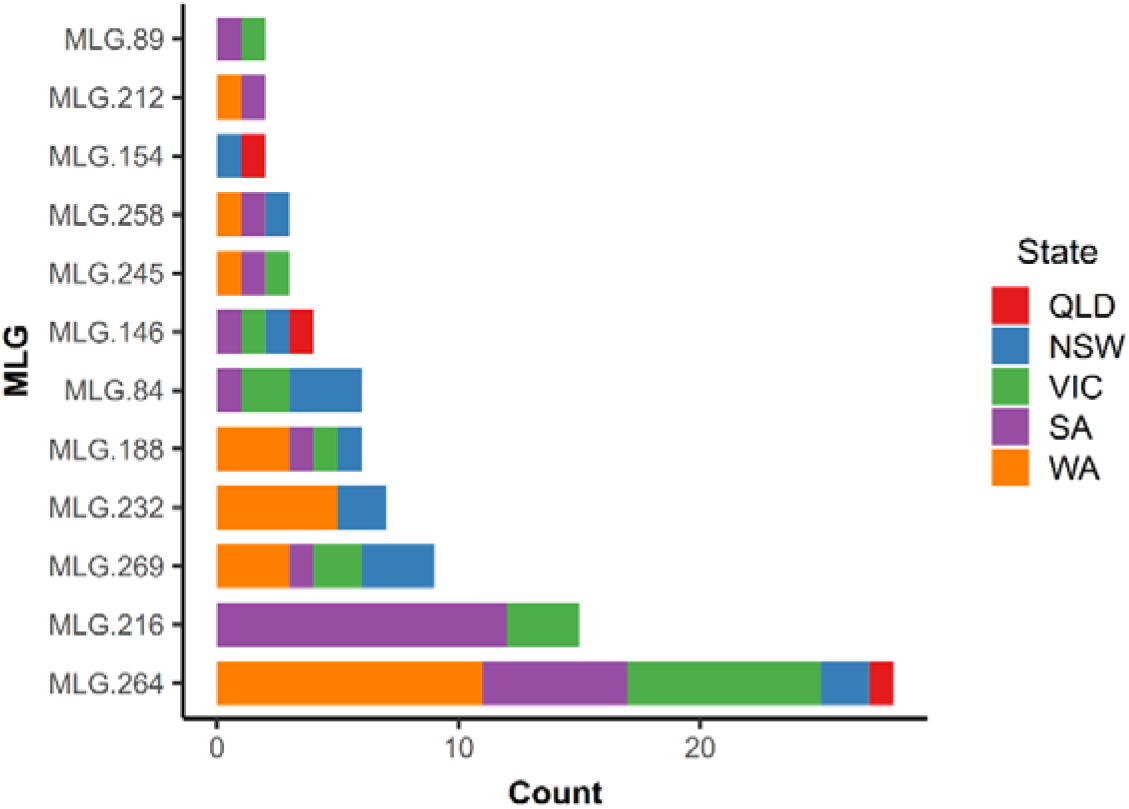
Multilocus genotypes of *Ascochyta rabiei* isolates sourced from multiple states in Australia, based on genome-wide SNP data.

### Association between *A. rabiei* genotypes and pathogenicity

The same MSN diagram of the MLG haplotypes, coloured to highlight the pathogenicity groups of the isolates, showed no distinct pattern of closely related highly pathogenic isolates, except for two diverged groups of isolates consisting mostly of isolates of pathogenicity groups 3-5 (red arrows in Figure 5).

**Figure 5.**
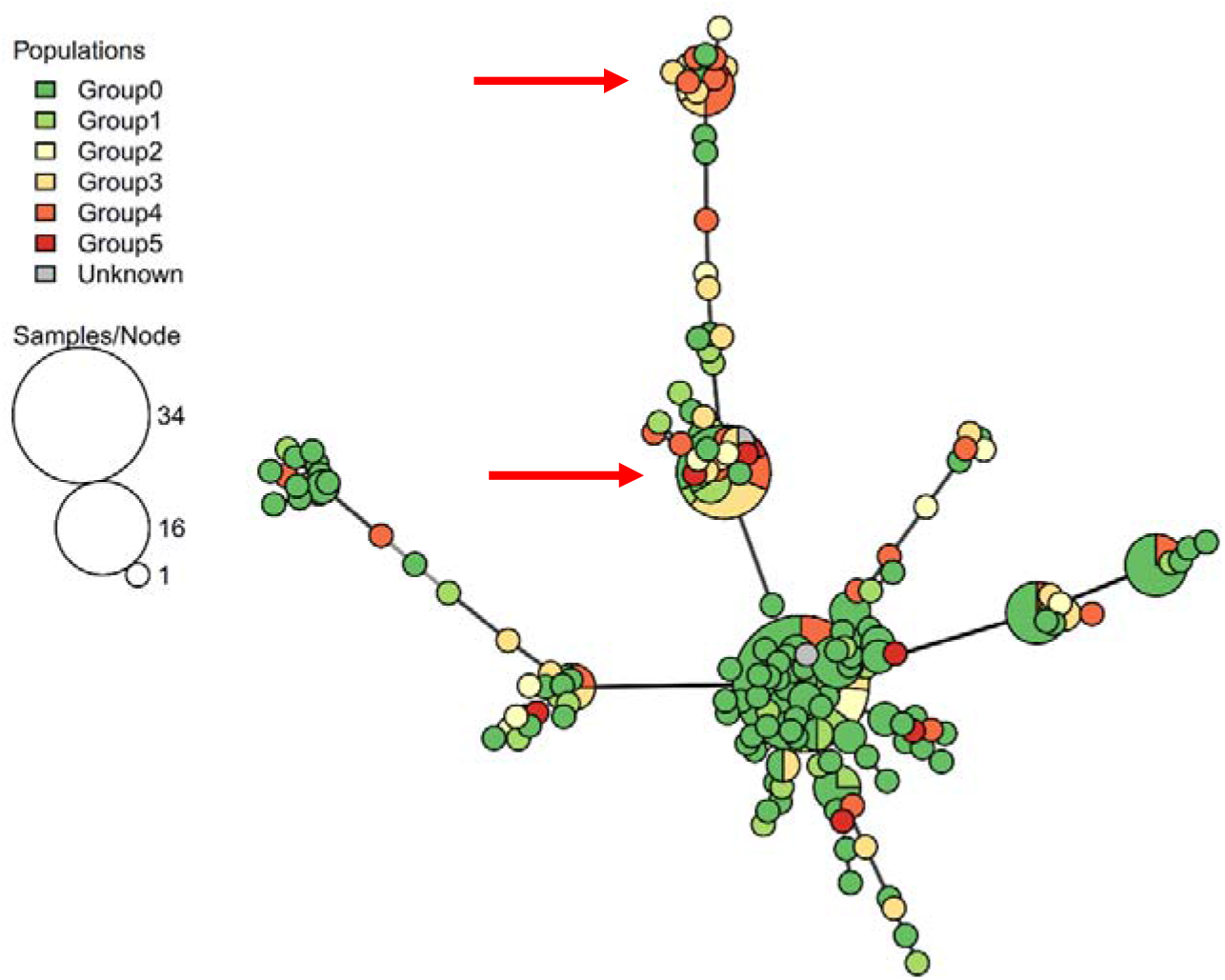
Minimum spanning network diagram of Australian *Ascochyta rabiei* multilocus genotypes by pathogenicity group. Where node size represents the number of isolates identified in each MLG, and the length of the branches between the nodes represents the genetic distance. Node colours represent the pathogenicity group of the isolates in the MLG. Arrows highlight nodes/clusters with high proportions of highly pathogenic isolates (PG ≥ 3).

Visualising the pairwise genetic distances between the Australian isolates as a clustered heatmap and dendrogram (Figure 6) provided additional information to better interpret the level of relatedness between isolates and association with their state of origin and pathogenicity level.

**Figure 6.**
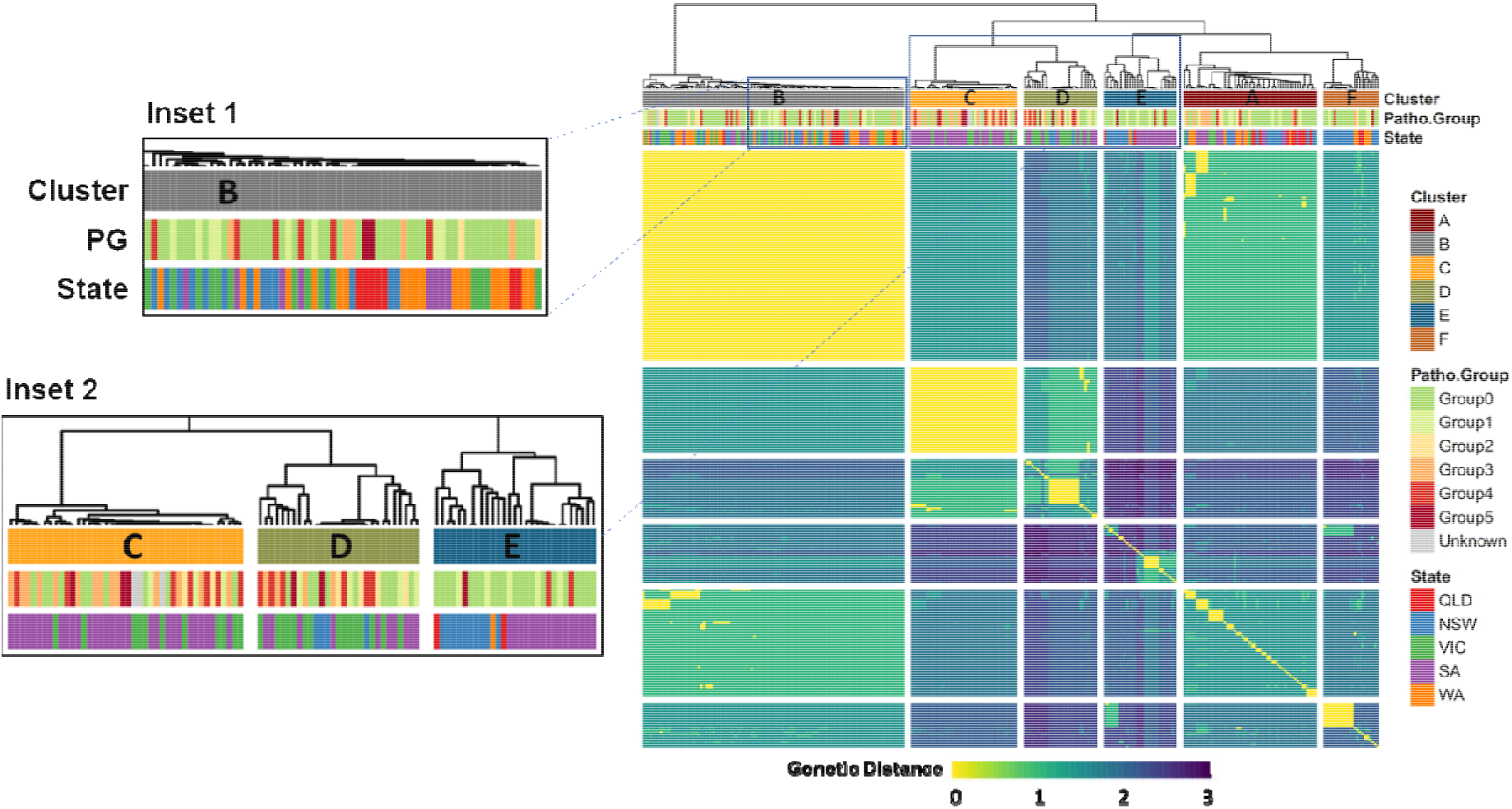
Genetic distance-based clustering of *Ascochyta rabiei* isolates. The heatmap and dendrogram are based on pairwise Euclidean distances that were calculated from SNP allele frequencies obtained from DArTseq™ data. Gaps in the heatmap separate between isolates clustered within major branches (denoted clusters A-F and coloured below the dendrogram according to the legend), as determined by hierarchical cluster analysis using the *hclust* function in R (default parameters used). Additional colour bars on top of the heatmap represent the Pathogenicity Group classification and State of origin of each isolate. Inset 1 shows a magnified view of genetically similar, but phenotypically diverse isolates in cluster B; Inset 2 shows a comparison between clusters C, D and E.

Overall, the hierarchical clustering that was applied was in agreement with the MLG determination, with just 2% of the MLGs including isolates from multiple clusters. MLGs 212, 232 and 245 included isolates from both clusters A and B, while MLG.174 comprised one isolate from cluster A and another from cluster F (Supplementary Table S1). The large MLG (MLG.264) and its closely surrounding relatives, seen at the centre of Figure 3 and in Figure 4, are clearly identified as cluster B in the heatmap and dendrogram in Figure 6. This cluster represents a range of isolates from several growing regions, years and pathogenicity groups (inset 1 in Figure 6), although they were almost genetically identical (clonal) according to their MLG assignment.

In contrast, other clusters showed strong association with these descriptors. For example, cluster **E** (bottom inset in Figure 6) comprised a distinct set of isolates originating from SA in 2013, along with NSW-originated isolates. Most of these Cluster E isolates (~90%) were of low pathogenicity (Table 4). Of further interest, Cluster **C** comprised mostly high pathogenicity isolates, all from SA and VIC (inset 2 in Figure 6). Based on the odds ratio, an isolate clustered within this group was at least ten times more likely to be classified as highly pathogenic (pathogenicity groups 3-5) than an isolate from clusters E or F, and at least six times more likely than an isolate from clusters A or B (Table 4 and Figure 6). A similar observation was made in the left branch of cluster D, although the isolates in this branch were less uniform genetically and the cluster included a branch of low-pathogenicity isolates (inset 2 in Figure 6).

**Table 4.** Proportions of virulence classification of *Ascochyta rabiei* isolates in dendrogram clusters.

| Cluster | High<sup>1</sup> | Low<sup>1</sup> | Odds Ratio<sup>2</sup> | Significance (*p*-value) |
| --- | --- | --- | --- | --- |
| A | 18.9% | 81.1% | C/A = 6.88 | 0.000877 (***) |
| B | 19.4% | 80.6% | C/B = 6.64 | 6.73E-05 (***) |
| C | **61.5%** | 38.5% | C/C = 1 | (ns) |
| D | 41.4% | 58.6% | C/D = 2.26 | 0.575 (ns) |
| E | 10.3% | **89.7%** | C/E = 13.87 | 0.002061 (**) |
| F | 13.6% | 86.4% | C/F = 10.13 | 0.012726 (*) |

<sup>1</sup>Isolates were classified as “Low” or “High” pathogenicity if they were classified in pathogenicity groups 0-2 or 3-5, respectively. Highest and lowest proportions are indicated in bold font.
<sup>2</sup>Differences in proportions of “Low” and “High” pathogenicity isolates between clusters were determined by a generalized linear mixed-effect model and are presented as the odds ratio (OR) compared to cluster **C**, along with their statistical significance as validated by the Tukey method.

GWAS analysis, utilising the same SNP set derived from the DArTseq™ data, revealed two SNP allelic variants, 44700443-64-A/G and 44702311-19-C/A, that were significantly associated with pathogenicity (). These markers were annotated to their genomic coordinates and were located on contigs 06 and 20 of the *A. rabiei* ArME14 reference genome (17) (NCBI accessions RYYQ01000007.1 and RYYQ010000021.1, respectively, see Table 5).

**Table 5.** SNPs associated with pathogenicity levels in *Ascochyta rabiei*

| SNP Name | Contig | Position | p-value | q-value |
| --- | --- | --- | --- | --- |
| 44700443-64-A/G | RYYQ01000021.1_ctg20 | 801717 | 6.5431E<sup>-10</sup> | 9.16034E<sup>-09</sup> |
| 44702311-19-C/A | RYYQ01000007.1_ctg06 | 1292348 | 5.54659E<sup>-05</sup> | 0.000388261 |

## Discussion

In this study, a genotyping-by-sequencing (GBS) approach was undertaken to assess the molecular genetic structure within the Australian *A. rabiei* population. Also, a select isolate core collection was interrogated to seek for potential correlation between isolate genotype and ability to cause significant disease severity on a differential host set.

### Population Structure and Diversity

With 212 polymorphic markers and 185 MLGs, the GBS approach offered a high-resolution set of markers for the analysis of genetic relatedness among the Australian isolates, and enabled comparison to the ICARDA *A. rabiei* isolates. This provided a more accurate measure of the genetic relatedness within the Australian collection than the seven microsatellite markers previously used by Mehmood et al. (11). The modest number of genome-wide polymorphic SNPs resulting from the DArTseq™ method in the current study, along with the level of clonality and genetic diversity metrics observed in the sampled population, demonstrated and confirmed the low level of genetic diversity previously reported for *A. rabiei* in Australia (11,12,46). Minor seasonal and regional variations in genetic diversity were observed, which may be a result of the environmental conditions and scale of cropping affecting the presence of the pathogen in the growing regions and hence in the isolate collection.

Inclusion of the two isolates from the ICARDA collection provided context to the homogeneity of the genotype data from the Australian isolates and confirmed that the DArTseq™ method detected many more SNPs than would be evident from the assessment of the Australian isolate data alone. The low number of polymorphic SNP loci in the Australian *A. rabiei* population is a consequence of genetic homogeneity rather than any inefficiency of the DArTseq™ method for genotyping fungi for population studies.

The genetic distances between Australian isolates, within and between collection locations (Figure 3 and Inset 2 in Figure 6) clearly showed that some isolates were highly related within a particular location, suggesting that they had evolved locally and potentially independently. In contrast, other MLGs represented isolates that may be considered as genetic clones, and these were sourced from multiple growing regions. This included MLG.146, MLG.188 and MLG.269, and MLG.264, which were sourced from all five states (Figure 4). MLG.264 was identified in cluster B in Figure 6 and was centrally positioned in Figures 2 and 3, connecting all other branches. MLG.264 comprised 28 isolates sourced from all growing regions, representing the “core” clonal lineage of *A. rabiei* in Australia, from which other lines diverged. A comparison between the MLG determination presented in this study and the SSR-based haplotypes assigned by Mehmood et al. [9], revealed six MLGs that contained isolates assigned to multiple SSR haplotypes. One might hypothesise that MLG.264, which was tightly clustered in the current analysis, equates to the most frequently observed haplotype ARH01, observed from the SSR genotyping. However, the MLG.264 haplotype contained isolates previously assigned to five different SSR haplotypes. Furthermore, an examination of the isolates that were previously assigned to haplotype ARH01 revealed that they were mapped to 77 different DArTseq™ SNP-based MLGs, suggesting that the seven SSR loci previously used were indeed limited in their genotyping ability.

Conversely, 14 isolates that belonged to the same clonal lineage (MLG) were assigned to separate and distinct clusters. Among these, nine were collected in WA (64%), at a frequency significantly higher than represented in the overall collection (17%). This is puzzling considering the high clonality of the population in WA, but it may provide evidence that WA isolates are derived from a few isolates that were imported from other regions and established within relatively few founder events during the expansion of the industry, thus explaining their genetic relatedness across clusters.

The presence of isolates from the same clonal lineages in multiple growing regions may be explained either by 1) Limitation of the marker set resolution or 2) That the isolates were derived from the same clonal lineage and have been transported between regions. In total 185 MLGs were identified among the population isolates assessed, indicating the ability to differentiate close relatives. The latter hypothesis is further supported by the fact that no MLG clusters were found to comprise only isolates collected from WA or QLD, rather these were always found in association with isolates from other regions (Figure 3 and Figure 6). Considering the geographic isolation of the chickpea growing region of WA, more than 1,500 km from the nearest chickpea growing regions in SA (Figure 1), and the known natural dispersal distance of *A. rabiei* clonal spores (up to 100m), it is highly unlikely that they have dispersed between the regions without human assistance(47,48).

Furthermore, the higher level of genotype diversity observed among isolates collected in QLD (Table 3) and their spread into multiple distant MLGs, suggested that these isolates were “imported” from other growing regions, probably through sourcing of contaminated seeds or spread by cropping equipment brought into the region. The chickpea industries in both QLD and WA have expanded rapidly in recent years, peaking in a “goldrush” for chickpea production during the 2016 and 2017 growing seasons, with the grain prices reaching record highs and above US$700 per tonne. During the same period, seed was imported to QLD from various sources and a large proportion of seed was not certified as “clean seed”. In addition, crop rotation best practice was frequently not observed between consecutive seasons (KM, personal comm). The increase in cropping intensity and scale, along with lax cropping practices likely resulted in the additional selective pressure on the populations leading to the steep increase in frequency of highly pathogenic isolates, observed in QLD during the 2017 season (22).

### Pathogenicity of the Australian *A. rabiei* population

Despite the low genetic diversity observed and the reduced rate of chromosomal mutations expected by clonal propagation, *A. rabiei* isolates in Australia exhibited a wide pathogenic range and were able to cause varying levels of disease on an established differential host set used to characterise them (11,22). Previous studies (46) and unpublished work by our team (RF, RL, IB) verified the absence of the second mating-type sequence (MAT 1-1), found elsewhere in the world, in the Australian *A. rabiei* population. Thus, the Australian pathogen is likely to reproduce in a clonal way, creating a self-selecting population of fittest and most pathogenic isolates (11). These fit and better adapted highly pathogenic isolates appear to be occurring at an ever-increasing rate, leading to shorter lifespans of newly bred and implemented resistant cultivars (9,22).

The variant SNP markers that were identified as associated with pathogenicity group were annotated by their chromosomal coordinate and association with potential virulence genes, such as effector molecules, signalling and receptor molecules, and transcription factors. One of the pathogenicity-associated markers, 44700443-64-A/G on ArME14 contig 20, was co-located in a gene-rich region (NCBI Genebank accession KR139658.1), the solanapyrone biosynthesis gene cluster. This includes genes related to metabolism, such as O-methyltransferase, dehydrogenase, polyketide synthase, and transcription factors. Kim *et al*. (2015) showed that knockdown of sol4, a novel type of Zn(II)2Cys6 zinc cluster transcription factor found in this region led to complete loss of solanapyrone biosynthesis, however this did not impact growth, sporulation or virulence of *A. rabiei* (49). Further investigation of this gene cluster and the Tc1/Mariner-type transposable elements surrounding it may identify their role in the infection mechanism and possible link to important protein-altering variants. The second trait-associated genomic region indicated by marker 44702311 contained no genes that could be associated with pathogenicity.

Genotyping by sequencing (GBS) and particularly DArTseq™ has become a common practice for large-scale genotyping of agricultural crops, animals and other species, thanks to its affordability compared with whole genome sequencing and SNP-chip assays and its focus on “functional” regions of the genome (26,50–52). GBS produces a set of markers throughout the genome, that is then used for population genetic analysis, genetic mapping, genome-wide association studies and other applications (53–55). Despite its advantages, a known caveat of GBS is the high rate of missing data, caused by the random representation of loci in each sample, which limits the number of comparable markers present in all samples, leading to sample and locus dropout (56).

In the case of the highly clonal Australian *A. rabiei* population, DArTseq™ resulted in just 212 informative SN Ps. This raises the question of whether the overall low population diversity and high clonality levels observed, particularly in clusters C and B, were a result of limited genotyping resolution or indicated a truly low level of genetic diversity. Previous studies have shown that a similar number of SNP markers spread over a small genome (41 Mbp for *A. rabiei* (17)) was sufficient to accurately determine population genetic structure, as was demonstrated for the fungal pathogen *Phytophthora rubi* (57).

To determine if the number of markers is sufficient to genetically distinguish between clonal lineages of A. *rabiei* and to obtain accurate marker-phenotype association to identify causative mutations in key genes, a higher marker density and coverage is required. The small genome size of A. *rabiei* and the continuous improvements in sequencing platforms along with reduction in sequencing prices are favourable to obtain this. In fact, a whole-genome-sequencing (WGS) or high-depth GBS approach to obtain high density sets of markers from large sample sizes may well be within reach in coming years. This will enable genome-wide association studies to identify specific alleles and loci correlated with pathogenicity levels, which could assist in resistance chickpea breeding and the development of novel fungicides. A similar approach identified novel candidate genes for pathogenicity in the wheat fungal pathogen *Fusarium graminearum* using approximately 30,000 SNPs spread throughout its 36 Mbp genome (58).

The set of SNP markers used in this study identified clonal isolates that were separated by just a few allelic variants, but displayed a wide phenotypic diversity. This may have indicated that the marker resolution was not sufficient to identify markers directly associated with pathogenicity and suggested that an isolate’s pathogenicity was not necessarily determined by a SNP variant. Other mechanisms that may alter pathogenicity may include structural factors, such as variation in copy number of a virulence gene, rearrangement of genes in the genome, duplication and translocation of transposable elements, effector repertoire and gene expression modifications, as well as epigenetic effects caused by specific host-pathogen interactions (59,60). Identifying these variations may require other genomic approaches, such as pangenome comparison incorporating long-read sequencing and transcriptomic approaches, including mRNA and short RNA sequencing (61,62). Application of these approaches in future studies will provide crucial evidence of genomic variations that are directly related to both pathogenicity and molecular interactions with the host resistance genes. The discovery of genetic cluster C, and to a lesser extent cluster D, that are indicative to a growing region and high pathogenicity, is encouraging in the search for distinct pathotype markers and provides a first step towards developing a molecular pathotype diagnostics method.

### Conclusions

This research aimed to elucidate the genetic structure of *A. rabiei* across the main chickpea growing regions of Australia and to determine any correlation of genotype with pathogenicity phenotype. To the best of our knowledge, this is the first published study that utilised a GBS approach and SNP markers to infer the genetic structure of an *A. rabiei* population, thus setting the foundation for ongoing investigation into specific genomic features of the pathogen that may assist Australian scientists, breeders, farmers and government agencies in managing and reducing the chickpea Ascochyta blight risk.

Although a direct genomic correlation to pathogenicity was not identified in the current study, an interesting pattern emerged suggesting that *A. rabiei* exhibits local micro-evolution within the broader growing region, likely driven by a founder effect, which has implications for local disease management and control of seed and farming equipment movements between the regions. The discovery of genetic clusters associated with highly pathogenic isolates holds a promise to the potential development of genetic markers that will enable early diagnostics of highly pathogenic *A. rabiei* isolates. This is crucial for accurate and informed disease management strategies.

These findings and further advances in the understanding of the pathogen populations will lead to the development of improved disease management strategies to allow sustainable production of chickpea as a source of food and income into the future.

## Author statements

### Authors and contributors

IB contributed to the conceptualisation of the research, performed the analysis, visualisation and curation of the data, and prepared the original draft of this work. PTS developed the methodology and performed the bioassays, sourced and single-spored the isolates and curated the isolate database. RCL and LMFC cultured the isolates and performed the DNA extractions. RCL also contributed to the data analysis and the writing of the manuscript draft. JD, KH and KM assisted in sourcing the isolates and provided useful input on the research design. RF supervised, conceptualised, secured the funding and provided project administration for the research. All authors reviewed, edited and provided valuable feedback during the preparation of this manuscript.

## Acknowledgements

*A. rabiei* isolates from the ICARDA collection were kindly provided by Prof. Diego Rubiales (Institute for Sustainable Agriculture, Cordoba, Spain). Australian isolates were kindly collected, processed and delivered by Merrill Ryan (QLD Department of Agriculture and Fisheries), Kurt Lindbeck and Gail Chiplin (NSW Department of Primary Industries), Jason Brand and Josh Fanning (Agriculture Victoria), and Marzena Krysinska-Kaczmarek (South Australian Research and Development Institute).

## Supplementary Files

**Figure S1.**
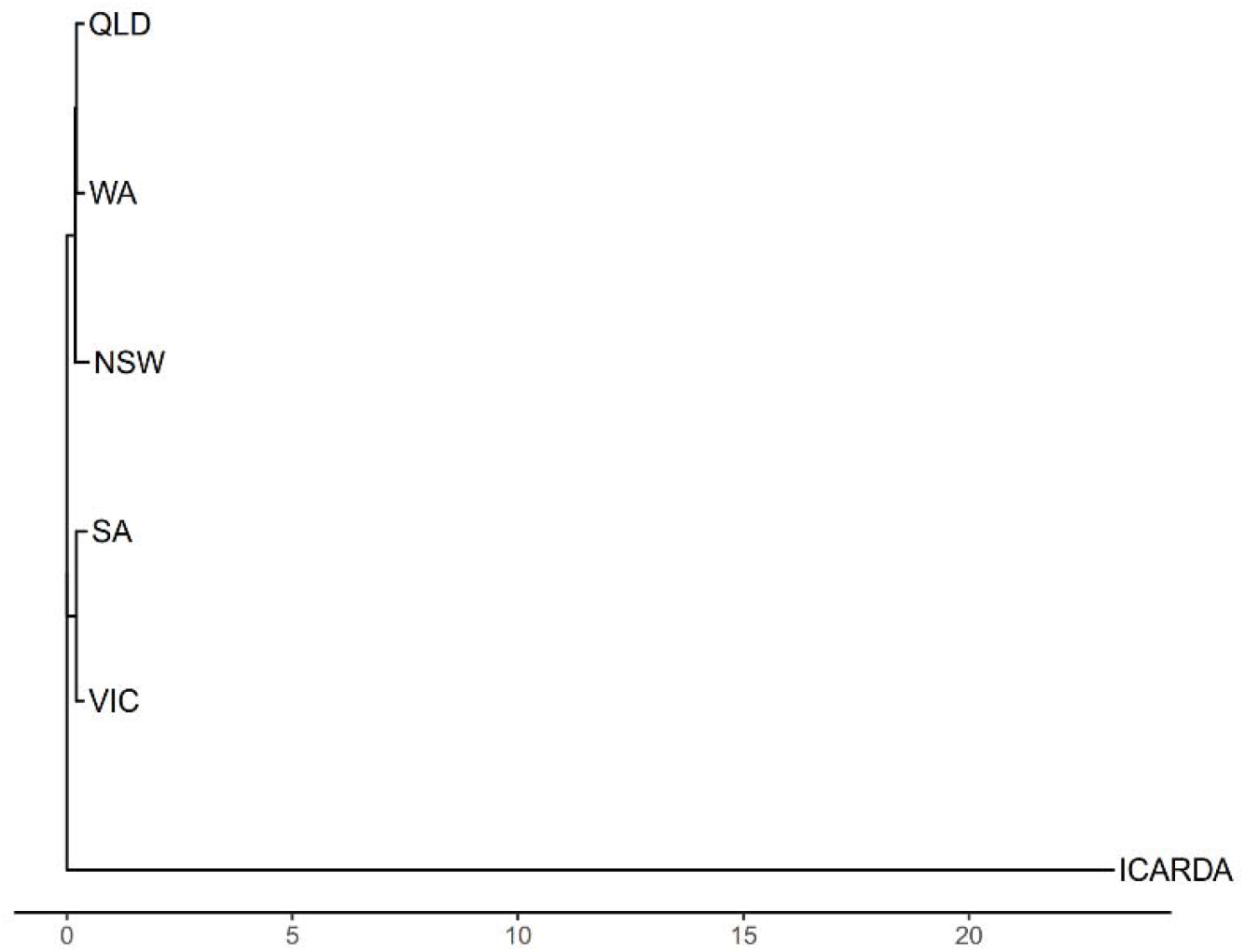
Neighbor-joining phylogenetic tree of *Ascochyta rabiei* populations

**Table S1.**
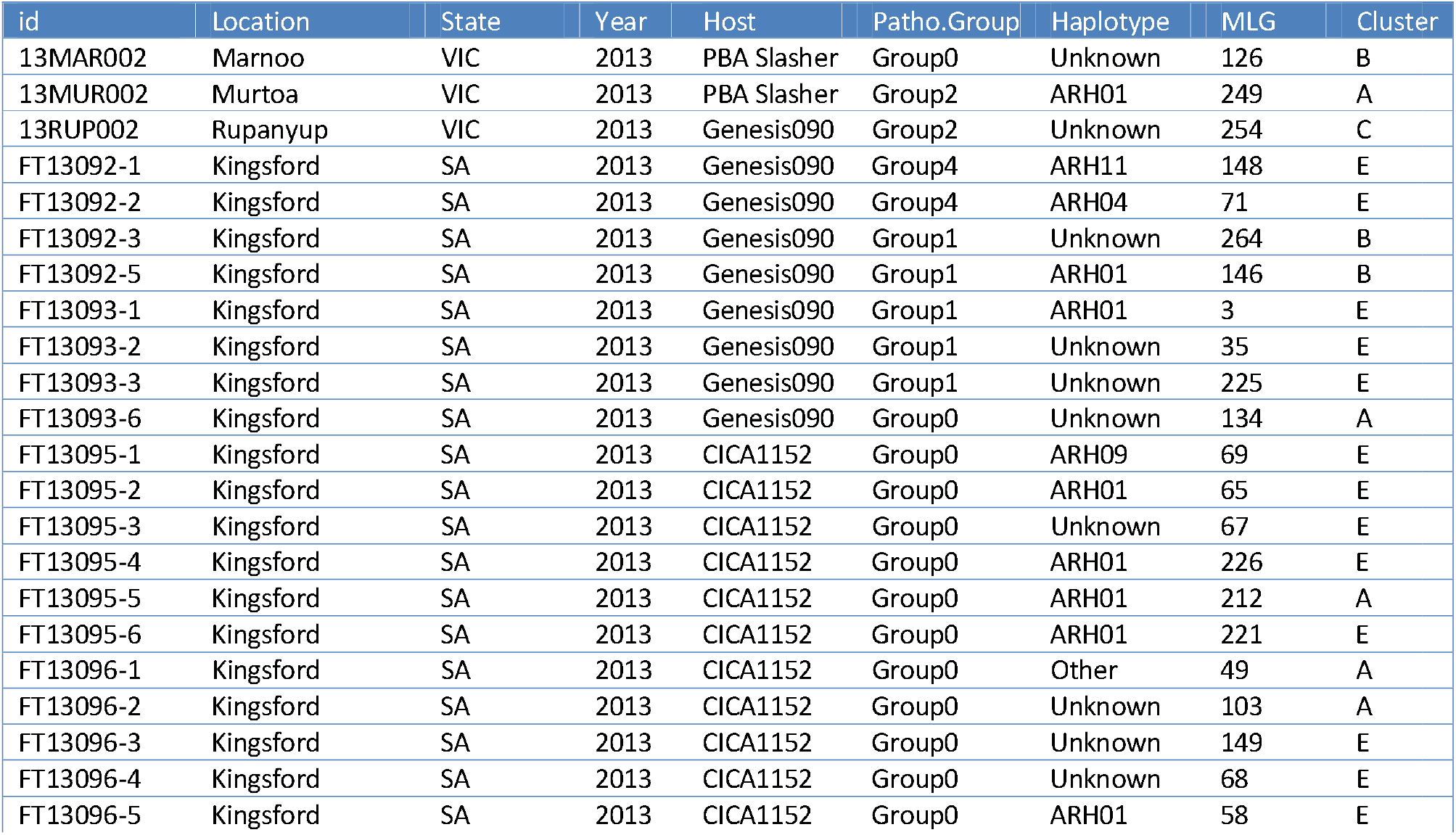
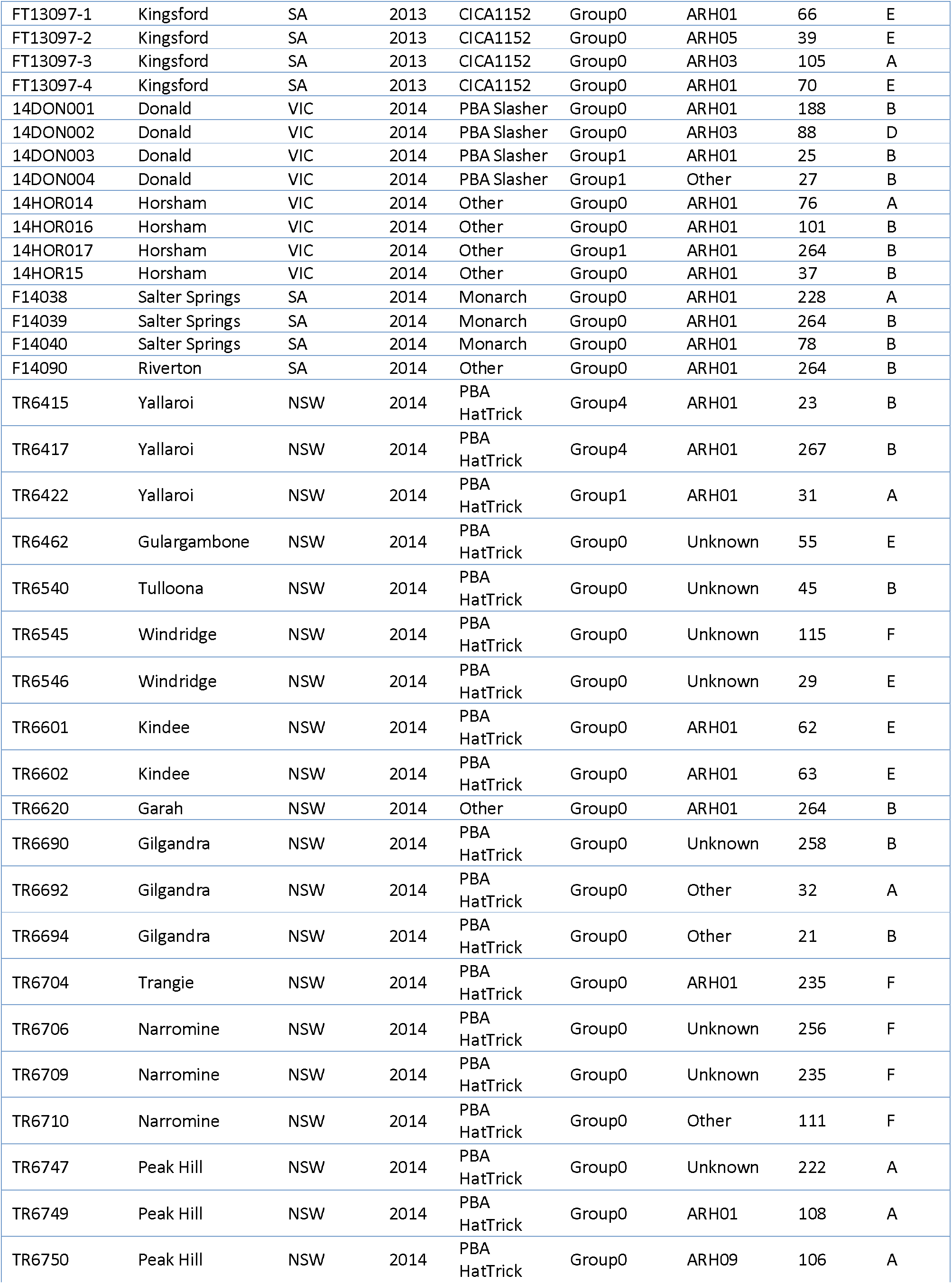
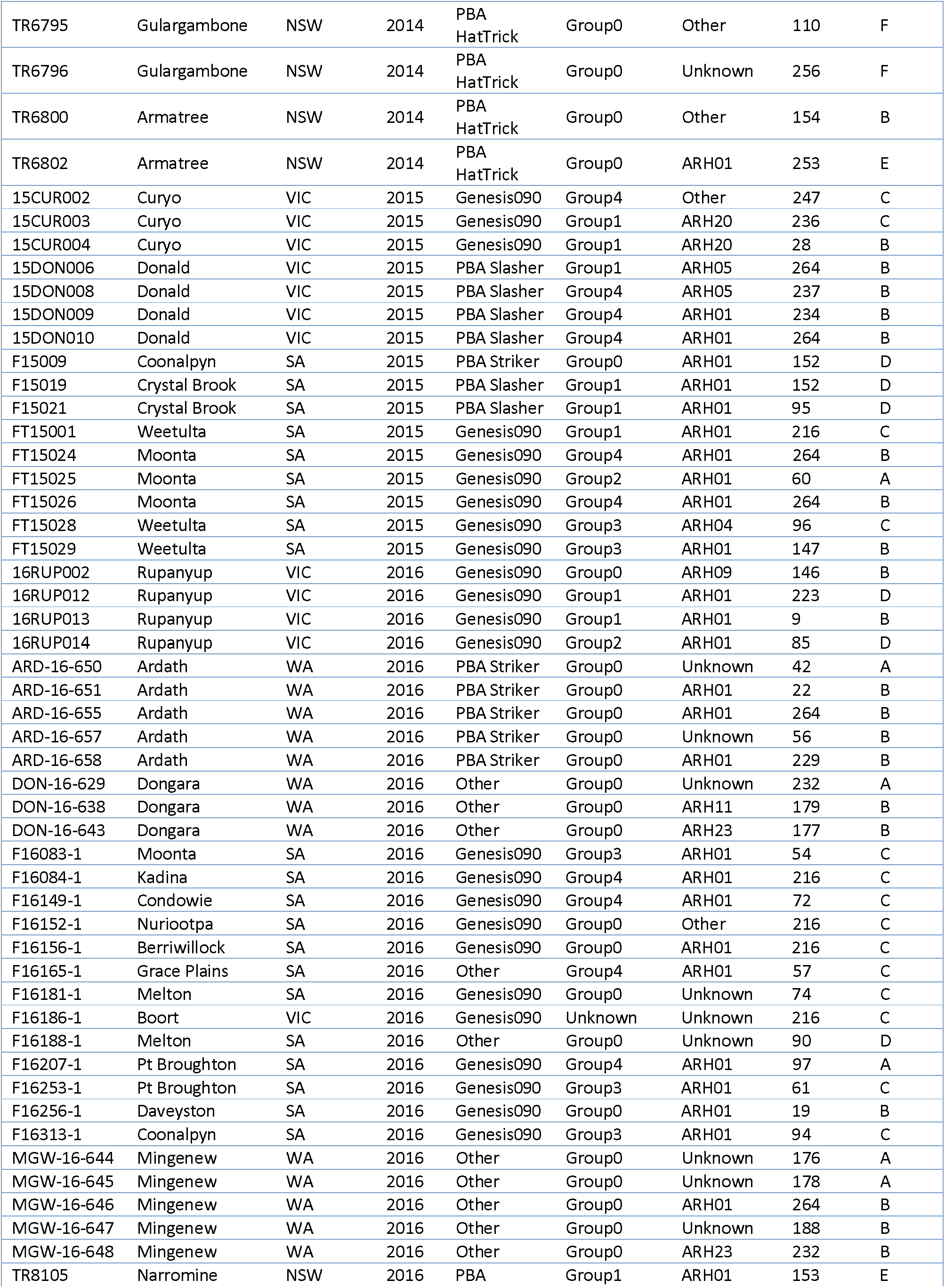
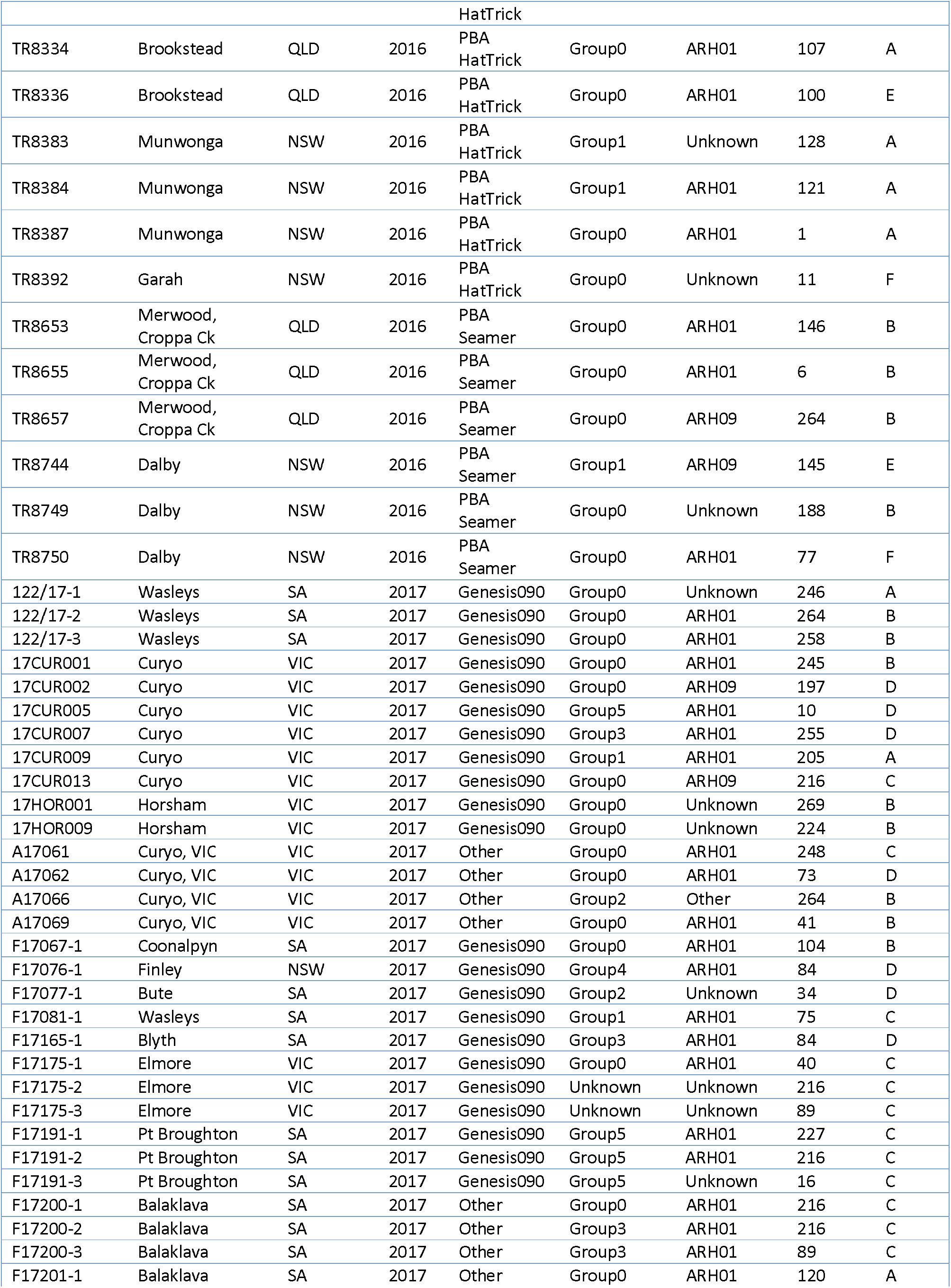
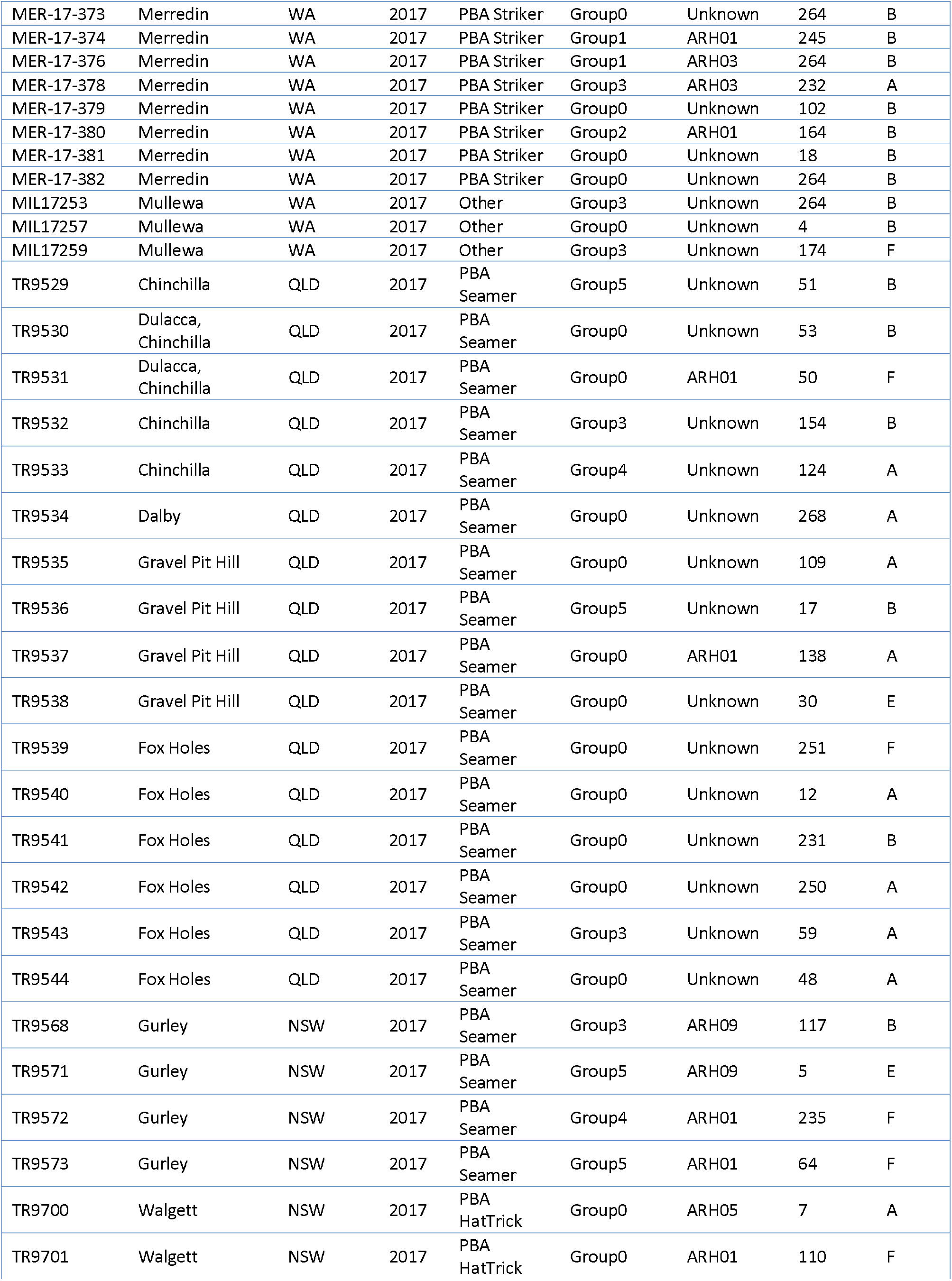
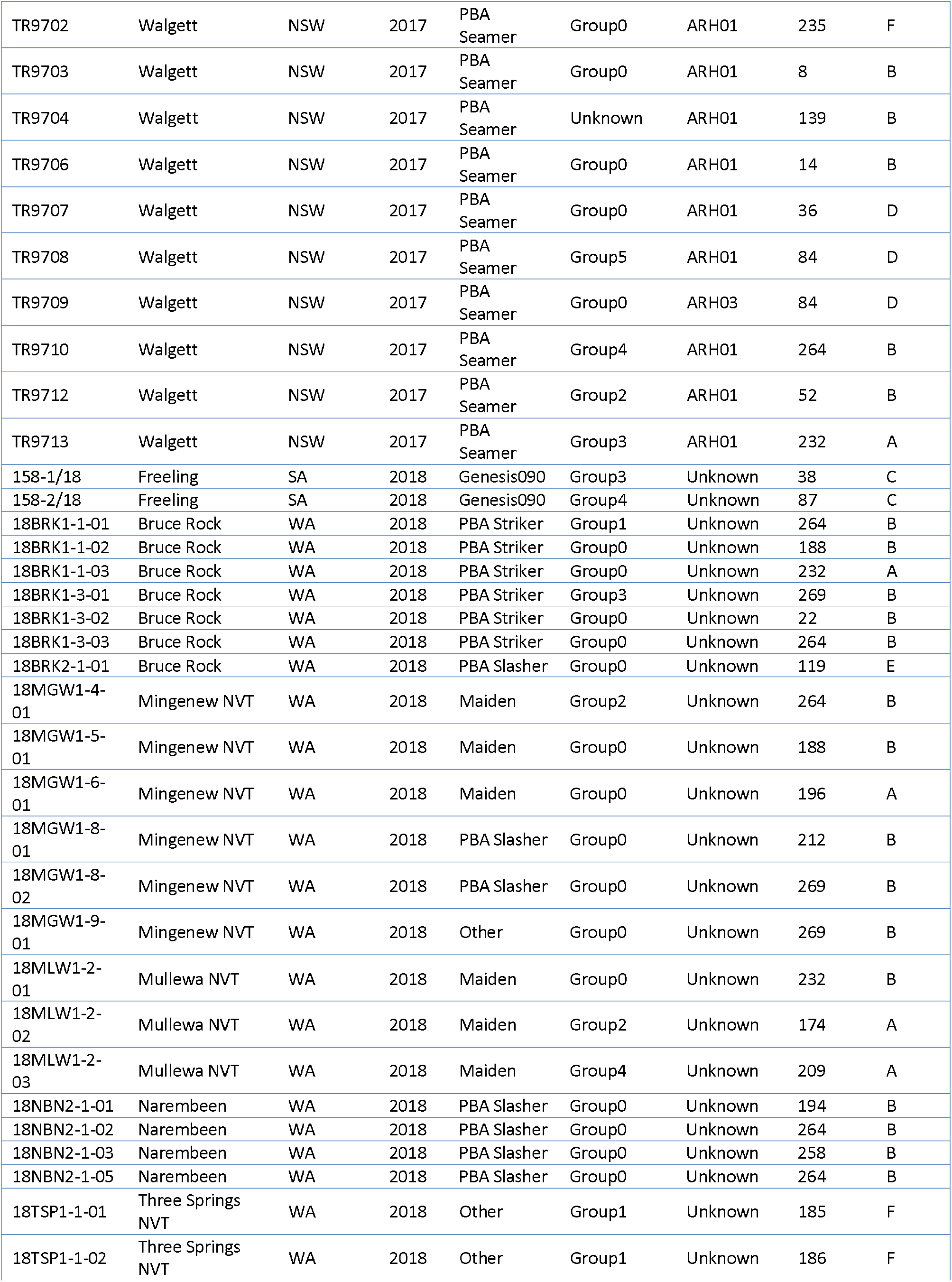
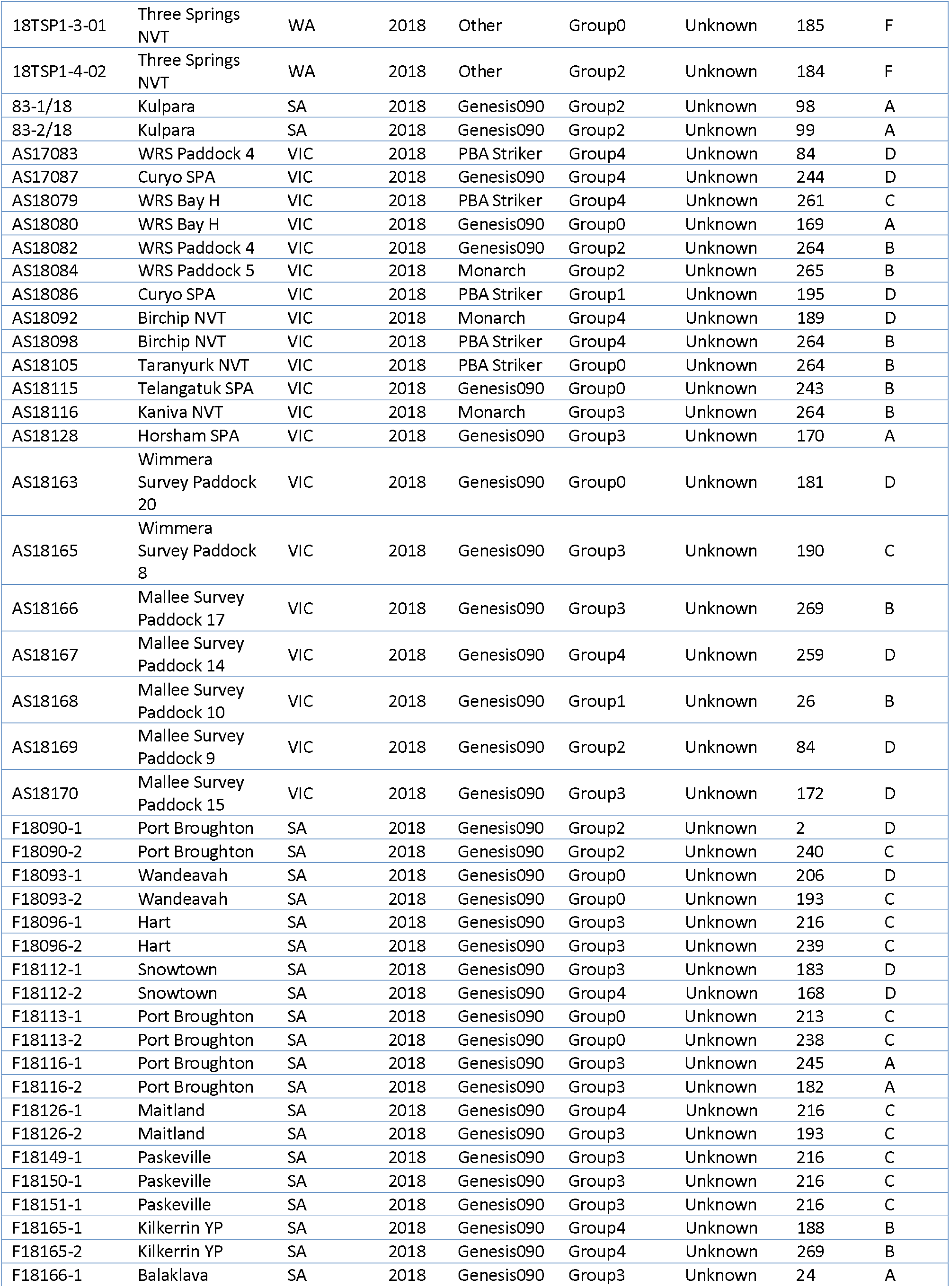
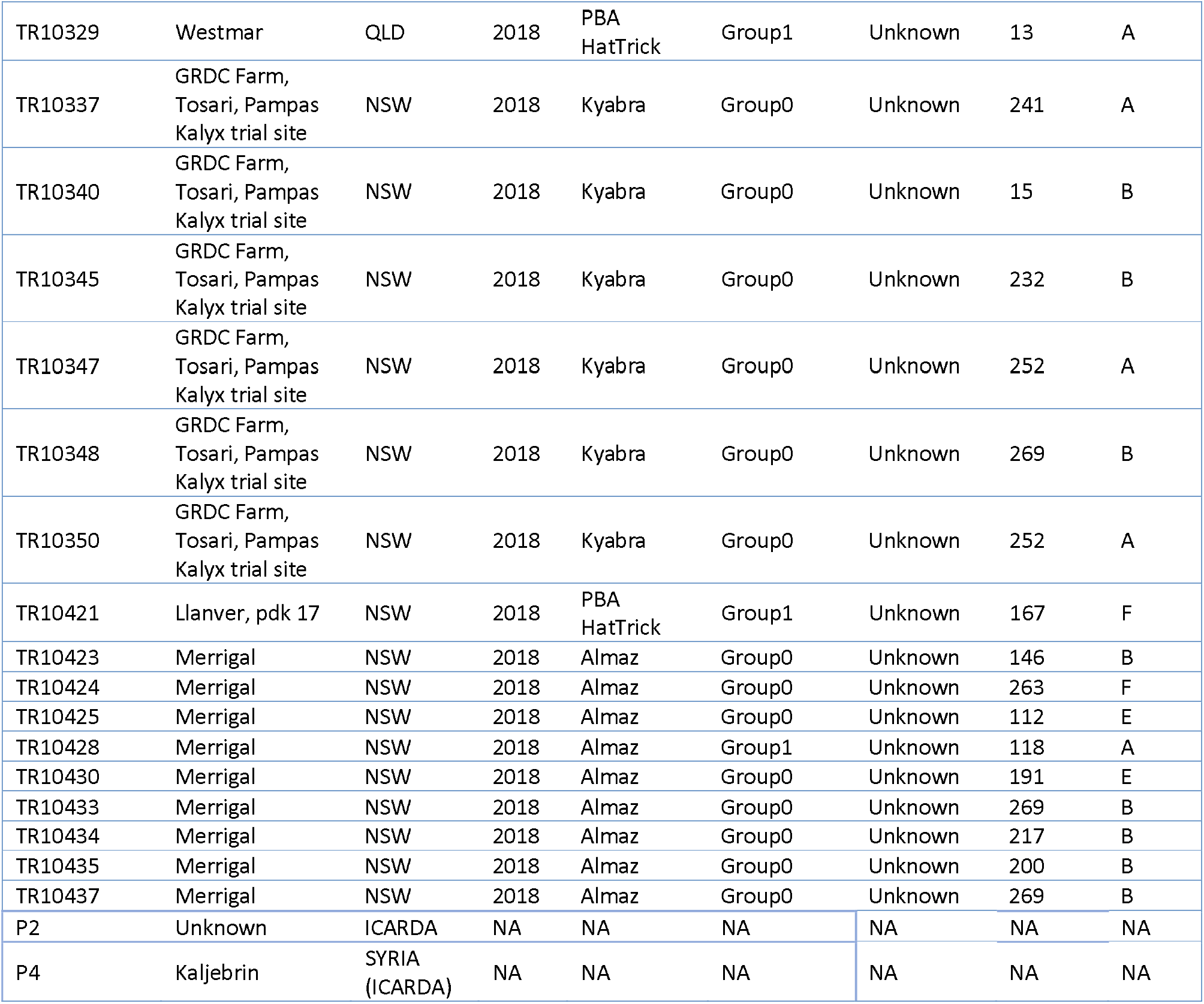
Details of *Ascochyta rabiei* isolates used in this study

